# Target-site Dynamics Explain a Large Share of Apparent MicroRNA Differential Expression

**DOI:** 10.1101/2025.09.29.679194

**Authors:** Mert Cihan, Piyush More, Maximilian Sprang, Federico Marini, Miguel A Andrade-Navarro

## Abstract

MicroRNA (miRNA) abundance reflects a dynamic balance between biogenesis, target engagement, and decay, yet differential expression analyses typically ignore changes in target-site availability driven by alternative polyadenylation (APA). We introduce MIRNAPEX, an expression-stratification-based machine learning framework that quantifies miRNA regulatory effect sizes from RNA-seq data by integrating target-gene expression with 3′UTR isoform usage to infer effective binding-site dosage. Using pan-cancer training sets, we train models that learn relationships between transcriptomic features and miRNA log-fold changes, with APA patterns providing predictive information beyond gene expression alone. When applied to knockdowns of core APA regulators, MIRNAPEX captured widespread 3′UTR shortening and accurately anticipated miRNA-specific shifts whose direction and magnitude mirrored APA-driven changes in binding-site availability. Analysis of target-directed miRNA degradation interactions further showed that loss of distal decay-trigger sites coincides with increased miRNA abundance, consistent with reduced degradation. Together, these findings demonstrate that apparent miRNA differential expression can arise from dynamic target-site landscapes rather than altered miRNA transcription, and that neglecting this dimension can lead to misestimation of regulatory effect sizes.

## Introduction

MicroRNAs (miRNAs) are short (∼22-nt) non-coding RNAs that post-transcriptionally regulate gene expression by binding to partially complementary sites in the 3′ untranslated regions (3’UTRs) of target genes [1]. By doing so, miRNAs fine-tune developmental programs, buffer cellular stress responses, and, if dysregulated, contribute to diverse disease phenotypes, including metabolic disorders, cancer, and neurogenerative diseases [2–5]. Considering the essential role of miRNAs in maintaining normal physiology and their potential as predictive biomarkers [6], accurate quantification of their expression level is crucial. In comparative transcriptomic analyses, differences in miRNA expression between conditions are commonly interpreted as indicators of altered post-transcriptional regulation and a reflection of broader regulatory state changes [7,8]. However, the steady-state level of a mature miRNA reflects a moving balance of three broad processes that determine the cellular abundance of each miRNA species. First, biogenesis, which includes transcription of the primary transcript, Microprocessor cleavage, nuclear export, Dicer processing and loading of Argonaute (AGO) proteins to form RNA-induced silencing complexes (RISCs), sets the potential pool of mature miRNAs [2,9]. Second, target engagement redistributes miRNA–RISC complexes across the transcriptome and determines how strongly a given miRNA can repress its targets in a particular cellular state [3,10,11]. Third, decay pathways remove mature miRNAs, with general turnover mechanisms such as tailing and trimming followed by exonucleolytic decay controlling their half-life [12,13].

In addition, target-directed miRNA degradation (TDMD) is a process in which binding to a highly complementary target transcript actively triggers destabilization and decay of the miRNA itself, thereby accelerating its decay. For instance, systematic AGO-CLASH analyses have revealed numerous endogenous transcripts that act as TDMD triggers, indicating that target-directed decay of miRNAs is more prevalent than previously recognized [14–16]. Because biogenesis, target engagement, and decay, including TDMD, act simultaneously and dynamically, observed changes in miRNA abundance may not directly report transcriptional output but can reflect shifts in target availability and turnover.

Accordingly, several studies have estimated miRNA activity from properties of their targets, showing that target gene abundance and site affinity predict miRNA levels, AGO binding, and competition effects [17–21]. Intuitively, target expression sets the demand placed on a miRNA, such that abundant, site-rich targets increase demand, whereas depletion of those targets reduces it [19,22,23]. However, most target-centric approaches treat each gene as if it had a single, fixed 3’UTR, overlooking the widespread phenomenon of alternative polyadenylation (APA) [24]. APA leads to the generation of transcript isoforms with distinct 3′UTR lengths and can therefore add or remove canonical miRNA binding sites as well as highly complementary TDMD trigger sites, dynamically altering the effective binding-site dosage available to each miRNA. Indeed, more than half of human genes utilize APA to generate alternative 3′UTR isoforms, meaning that dynamic 3′UTR length changes broadly modulate available miRNA binding sites and thus influence post-transcriptional regulation [24–27]. Since APA is a widespread mechanism that controls 3′UTR length and miRNA‐site availability, perturbation experiments of core APA factors (such as CFIm25, CFIm68 and CPSF6) have shown that knocking them down remodels thousands of 3′UTRs across the transcriptome [28,29]. Such APA-driven alterations in miRNA targeting have been shown to impact gene expression programs stem cell function and differentiation, and oncogenic transformation, among others, highlighting the crucial interplay between APA and miRNA regulation in fine-tuning cellular phenotypes [30,31].

Despite this, most DE analyses of miRNAs ignore dynamic changes in effective target-site dosage caused by APA and their impact on TDMD, creating a gap in how observed miRNA shifts are interpreted. We therefore hypothesize that APA-driven 3′UTR remodeling, by altering binding sites, systematically shapes mature miRNA levels such that effective target-site availability correlates with the effect size of miRNA regulation, typically quantified as log fold-change (logFC), between states and samples. Consequently, apparent miRNA DE often reflects target-site dynamics rather than altered miRNA transcription.

Motivated by this gap, we introduce MIRNAPEX, an expression-stratification-based interpretable machine learning (ML) framework that predicts miRNA logFC from RNA-seq by integrating target-gene expression with 3′UTR isoform usage to estimate effective binding-site dosage. Beyond prediction, we quantify the relative impact of APA variation on each miRNA and apply MIRNAPEX to APA-factor perturbation datasets to test whether global 3′UTR shortening produces predictable shifts in miRNA levels. We further examine curated TDMD trigger–miRNA pairs to see if loss of distal TDMD sites coincides with expected increased miRNA abundance [16]. Altogether, this approach shows how transcriptomic variation, including 3′UTR remodeling, shapes miRNA abundance, underscoring that miRNA DE and its estimated effect size should be interpreted in the context of dynamic target-site landscapes.

## Methods

### Data Collection

To train the ML models for predicting miRNA logFCs based on RNA sequencing data, we assembled datasets from The Cancer Genome Atlas (TCGA) [32]. We downloaded all available mRNA and miRNA quantification data from TCGA and cross-referenced these samples with the TC3A database [33], a resource that applies the DaPars algorithm to TCGA RNA-seq data to quantify APA patterns [34]. APA is represented by percentage of distal usage index (PDUI) values, that serve as a measure for distinguishing long and short 3′UTRs. PDUI values range between 0 and 1. For miRNA we obtained isoform-level quantification files and mapped them to mature miRNA entries using miRBase annotations [35].

The final dataset comprised 8460 samples with matched mRNA, miRNA, and APA profiles. Specifically, it includes TPM values for gene expression, mean RPM values for 2,000 mature miRNAs, and PDUI values for between 1058 and 11,266 genes per cancer type. Because APA usage is influenced by gene expression and biological context, the number of genes with valid PDUI values varies across cancer types. In total, the dataset spans 32 distinct TCGA cancer types and forms the basis for training the miRNA-specific ML models.

### Feature Engineering and Sample Definition

For each miRNA, putative target genes were obtained from the microT database [36] and ranked according to their gene-level microT interaction scores. To systematically evaluate the impact of feature set size, we constructed multiple input variants per miRNA by selecting the top 25, 50, 75, 100, 250, 500, 750, 1000, and 2000 highest-scoring target genes that are reported to have APA measurements. To generate training examples for each miRNA, we first randomly split all available samples into training (80%) and test (20%) sets. For model evaluation, the training set was further divided into five folds for cross-validation (CV). Within each fold, samples were stratified into high and low expression groups based on the expression level of the respective miRNA. Each sample from a specific fold was then randomly paired with a sample from the opposite expression group within the same fold. This strategy enabled the creation of diverse sample pairs representing varying expression differences while preventing data leakage between training and validation subsets. For each generated sample pair, we computed gene-level differential features for all target genes. Specifically, we calculated the logFC in mRNA expression between the two samples and the corresponding difference in PDUI values (ΔPDUI) for APA usage. To avoid undefined values due to zero expression, a correction of +1 was applied prior to logFC calculation. ΔPDUI values range between −1 and 1, reflecting relative changes in distal polyadenylation site usage. Features are computed in the same way for the test and full training sets (Figure 1).

**Figure 1.**
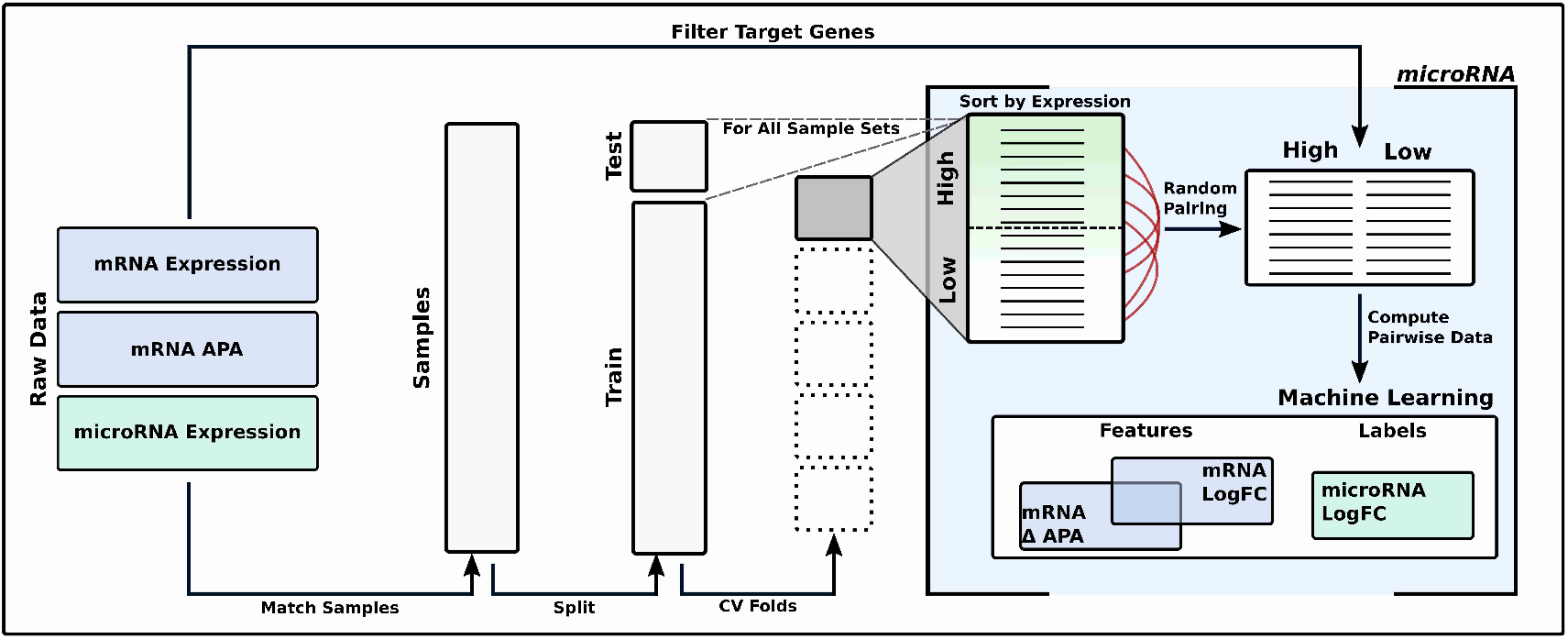
Workflow for training ML models to predict miRNA logFC. Matched TCGA samples containing mRNA expression, APA profiles, and miRNA expression are split into random training (80%) and test (20%) sets, with the training set further divided into cross-validation folds. Within each fold, samples are stratified by miRNA expression levels and randomly paired in both directions to prevent data leakage during training. For each miRNA, annotated target genes with valid APA measurements are selected, and differential features (mRNA logFC, ΔPDUI) are computed for each sample pair. The observed miRNA logFC serves as the prediction label, and the same feature computation is applied across folds, the test set, and final training set after hyperparameter tuning. A separate ML model is trained for each miRNA to capture the relationship between transcriptomic changes and miRNA expression dynamics.

### Training ML models to predict miRNA expression changes

We trained a range of ML algorithms to predict logFC values of mature miRNAs using the transcriptomic feature sets described above. To evaluate model performance under varying input conditions, we generated multiple training datasets based on different random pairings of samples and varying numbers of miRNA target genes (ranging from 25 to 2000). For each configuration, we trained both linear and non-linear regression models implemented in the scikit-learn library [37], including ordinary least squares (OLS), Lasso (LA), Ridge (RI), Elastic Net (EN), histogram-based gradient boosting regressor (HB), random forest regressor (RF) and multilayer perceptron (MLP). Hyperparameters for each model were optimized using CV within the training folds. For each miRNA, the model and hyperparameter combination that achieved the highest mean R^2^ across CVs was selected and retrained on the full training set to produce the final predictive model.

### The MIRNAPEX workflow

The resulting miRNA-specific ML models form the core of MIRNAPEX, enabling prediction of miRNA logFC values between two groups of RNA-seq samples. The MIRNAPEX pipeline automates the full process, starting from the raw FASTQ files. It integrates the GDC mRNA quantification pipeline [38] and DaPars-based APA analysis [33,34,39] to compute gene-level logFC and ΔPDUI values for predefined miRNA target genes. These features are then passed to pretrained miRNA-specific regression models to predict logFC values for 1165 miRNAs across any two user-defined sample groups (Supplementary Figure 1).

### APA perturbation

To test whether APA-driven changes in binding-site availability translate into shifts in mature miRNA levels, four RNA-seq comparisons of APA-regulator knockdowns with matched controls were analyzed. These perturbations remodel 3′UTRs, altering binding-site dosage. MIRNAPEX was then applied to predict miRNA log-fold changes between perturbed and control samples, and concordance with APA-driven target-site changes was evaluated. The datasets involve CFIm25 knockdown in HCT116 (GSE158591) as CFIm25-KD-1 [40]; CPSF6 knockdown in HEP3B (GSE229281) as CPSF6-KD [41]; and the HEK293 experiments comprising an independent CFIm25 knockdown replicate and a CFIm68 knockdown (GSE179630) as CFIm25-KD-2 and CFIm68-KD [29], respectively. We validated mature miRNA expression levels in respective cell lines using DIANA-miTED [42] and annotated miRNA binding sites on target transcripts with predictions from the DIANA-microT [36].

To approximate transcriptional contributions to miRNA abundance, we derived intronic transcriptional proxy measurements for a subset of TDMD pairs in which the miRNA is encoded within an intron of the trigger or host transcript. These measurements capture changes in transcription at the pri-miRNA locus and were used to evaluate whether mature miRNA logFC could be explained by altered transcription.

## Results

### Transcriptomic Prediction of miRNA expression changes

miRNA logFC values between sample groups were predicted using features derived from their putative target genes. Two types of features were considered: gene-level logFC values, reflecting differential mRNA expression, and ΔPDUI values, capturing changes in APA patterns. Together, these measures serve as proxies for the relative abundance of miRNA binding sites within their target genes. To avoid reliance on a single modeling assumption, we compared several commonly used regression algorithms that differ in their treatment of high-dimensional feature spaces. These include linear models with L1 or L2 regularization, which emphasize feature selection or coefficient shrinkage, as well as non-linear models that capture complex relationships through ensemble or kernel-based approaches. To assess predictive performance, ML algorithms were built using feature sets of varying size, defined by ranked microT interaction scores (see Methods for details).

Linear models substantially outperformed non-linear approaches in predicting miRNA logFC values across feature set sizes (Figure 2A). EN achieved the highest mean R^2^, followed closely by LA and RI, while non-linear models such as RF, HB, and MLP performed worse. As a baseline algorithm, OLS exhibited a marked decline in performance once the feature set exceeded ∼250 genes, highlighting the importance of regularization in high-dimensional settings. These findings are consistent with prior observations that linear models are well suited for modeling miRNA expression dynamics [21].

**Figure 2.**
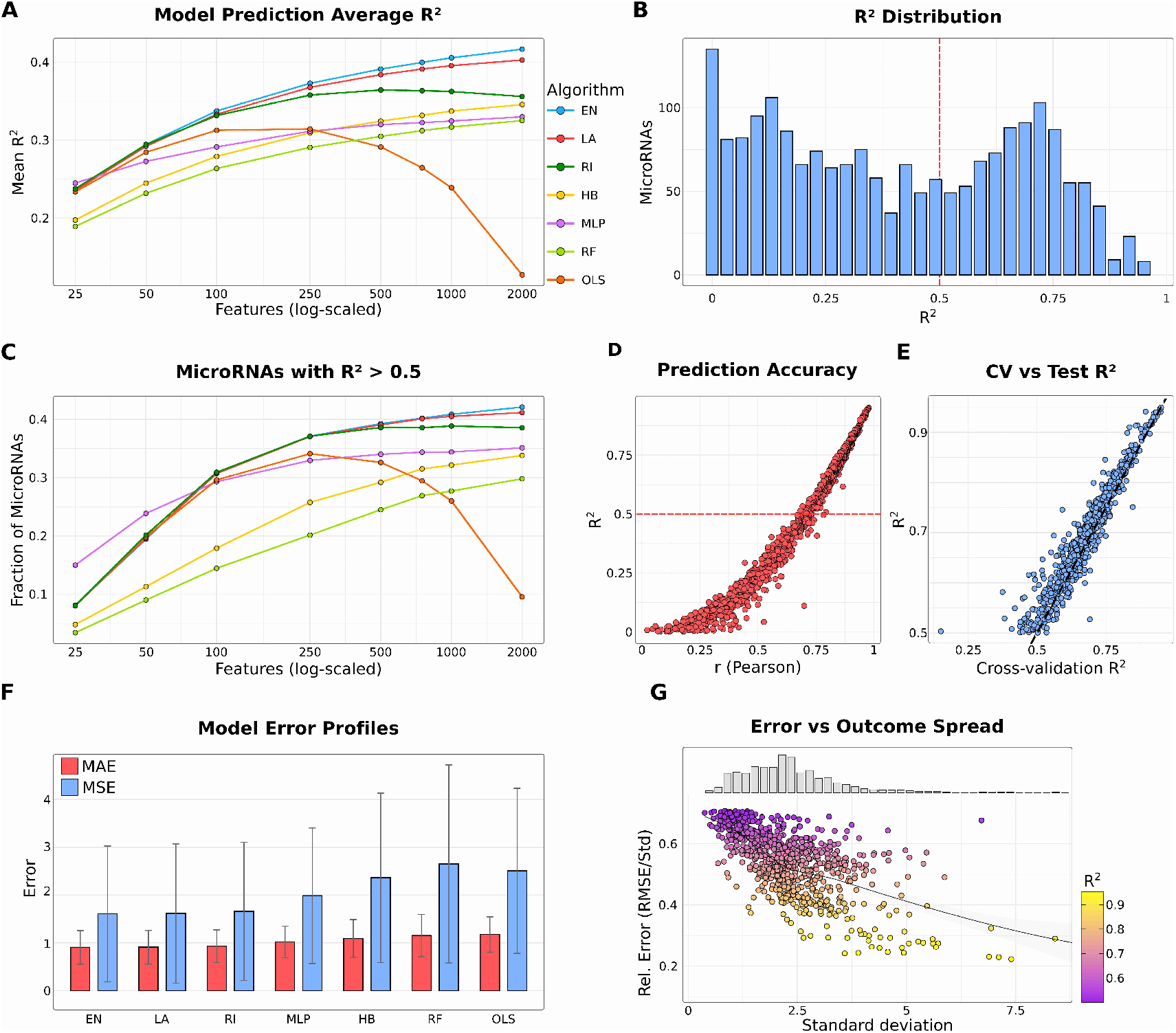
Prediction performance of ML models for miRNA logFC. (**A**) Line plot of mean R^2^ values across different ML algorithms as a function of the number of input features (tick marks represent logarithmically spaced values). Algorithms are abbreviated as follows: EN, elastic net; LA, lasso; RI, ridge regression; HB, HistGradientBoost; MLP, multilayer perceptron; RF, random forest; OLS, ordinary least squares. (**B**) Distribution of R^2^ values from EN trained with 1000 features across all evaluated miRNAs. (**C**) Fraction of miRNAs achieving R^2^>0.5 as a function of the number of input features (tick marks represent logarithmically spaced values). (**D**) Dot plot of Pearson correlation between predicted and observed logFC values versus miRNA-specific R^2^. (**E**) Comparison of cross-validation R^2^ against test set R^2^ across miRNAs. (**F**) Mean absolute error (MAE) and mean squared error (MSE) across algorithms (for 1000 features). (**G**) Relationship between the standard deviation of predicted logFC values and relative error (RMSE/standard deviation). The distribution of standard deviation values is shown as a histogram.

A feature set size of 1000 was selected as the optimal balance between predictive accuracy and interpretability (Supplementary Table 1). At this scale, EN achieved a mean R^2^ of 0.41 across all miRNAs, and the distribution of prediction accuracies showed that 817 miRNAs (41%) surpassed the R^2^ > 0.5 threshold, with a mean R^2^ of 0.69 for this subset (Figure 2B,C). To define highly predictable miRNAs (HP-miRNAs), we used a threshold of R^2^ > 0.5, corresponding to models that explain more than half of the observed variance. This cutoff aligns with the right-hand tail of the R^2^ distribution (Figure 2B). These HP-miRNAs were prioritized for downstream analyses. Across all miRNAs, the average Pearson correlation between predicted and observed logFC values was 0.62, while the correlation increased to 0.83 for HP-miRNAs (Figure 2D). Robustness of the models was further supported by the strong concordance between cross-validation and test set performance, with a Pearson correlation of 0.98 (Figure 2E), indicating minimal over- or underfitting.

To further benchmark model accuracy, prediction errors were compared across algorithms at the 1000-feature setting (Figure 2F). EN achieved the lowest mean absolute error (0.91) and mean squared error (1.60), further highlighting its robustness relative to the other methods. Moreover, analysis of relative error revealed that prediction error scaled proportionally to the variance of the observed logFC values and remained small relative to the standard deviation, particularly for HP-miRNAs (Figure 2G).

Together, these analyses demonstrate that effect size estimate of the miRNA expression regulation can be predicted with high accuracy and robustness from transcriptomic features.

### Expression- and APA-driven signals jointly shape miRNA prediction accuracy

The prediction of miRNA activity from transcriptomic data has traditionally been based on mRNA expression levels measured by RNA-seq or microarray platforms [43,44]. To investigate the added predictive value of 3′UTR patterns, we evaluated the role of APA. Specifically, we trained ML models for each miRNA using three different feature sets: expression-only, APA-only, and combined expression plus APA features.

Expression-only models achieved a mean R^2^ of 0.39 across all miRNAs, while APA-only models performed slightly lower with a mean R^2^ of 0.36. Importantly, the combined models improved performance to a mean R^2^ of 0.41, demonstrating that APA contributes complementary predictive information beyond gene expression alone (Figure 3A). Among the HP-miRNAs, there were 802 miRNAs for the expression-only and 617 for the APA-only models scoring with R^2^ > 0.5. Although the gain in overall prediction performance when using expression and APA features together is modest compared with using either feature set alone, this analysis demonstrates two important points. First, APA-only models perform comparably to expression-only models, indicating that a measure independent of gene expression quantification, namely 3′UTR length patterns, can predict differential miRNA behavior. Second, combining expression and APA features provides a unified framework to assess their relative and context-dependent contributions to miRNA regulation. To assess model performance on high-confidence miRNAs, we evaluated MIRNAPEX predictions for MirGeneDB-supported miRNA genes [45]. Across all MirGeneDB miRNA entries, the mean predictive accuracy was R^2^ of 0.61. At the gene level, allowing either mature arm to contribute, 364 of 506 MirGeneDB miRNA genes showed strong predictability (R^2^ > 0.5). Together, these results indicate that MIRNAPEX performance is strongest for high-confidence miRNA annotations and is not driven by low-confidence miRBase entries.

**Figure 3.**
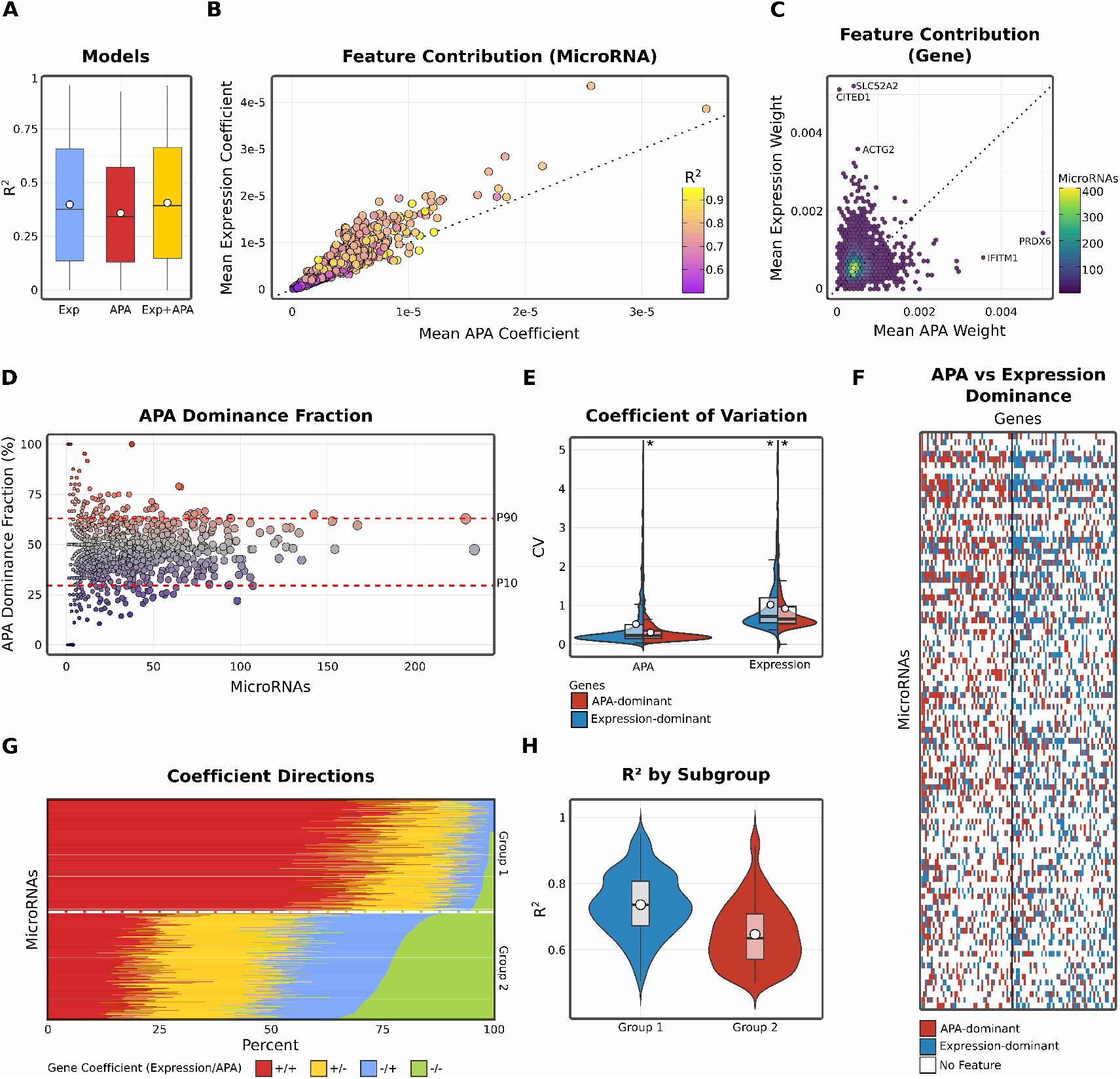
Contribution of APA and expression features to miRNA logFC prediction. (**A**) Comparison of predictive performance for models trained with APA-only, expression-only, or combined features. Boxplots show the distribution of R^2^ scores across miRNAs. (**B**) Scatter plot of average normalized absolute coefficients for APA versus expression features across miRNAs with R^2^>0.3. Each point represents one miRNA, colored by predictive performance (R^2^). (**C**) Hexbin plot of gene-level contributions, showing mean percentage weight of APA versus expression features across highly predictable miRNAs (R^2^ ≥ 0.5). Color scale denotes the number of miRNAs in which a given gene contributes. (**D**) Scatter plot of APA dominance fraction versus gene prevalence across all genes contributing to miRNAs with R^2^ ≥ 0.5. Each point represents one gene, with dashed lines marking the 10th and 90th percentile cutoffs used to define expression-dominant versus APA-dominant gene sets. (**E**) Coefficient of variation for APA versus expression features within the extreme APA-dominated and expression-dominated gene deciles. These measurements represent the raw variability from which APA and expression features are derived for model training. Stars denote outliers above the plotted range. (**F**) Heatmap of categorical dominance (APA versus expression) across all miRNAs with R^2^ ≥ 0.5 and the 100 most prevalent genes. Rows are ordered by decreasing miRNA R^2^, and columns by decreasing gene prevalence. White indicates that the gene was not a selected feature for that miRNA, blue indicates higher absolute expression coefficient, and red indicates higher absolute APA coefficient. (**G**) Stacked barplots of sign concordance between APA and expression coefficients across all genes for miRNAs with R^2^ ≥ 0.5. Colors denote the four possible sign combinations: both positive (++), APA positive with expression negative (+−), APA negative with expression positive (−+), and both negative (−−). miRNAs are stratified into two groups based on their composition, using a threshold of 10% (−−). (**H**) Comparison of R^2^ values between the two stratified groups.

To further investigate the predictive signal, we examined feature contributions from both the miRNA and target gene perspectives.

From the miRNA perspective, analysis of average coefficients confirmed that both expression- and APA-derived features contributed substantially to prediction accuracy, with no miRNA relying exclusively on a single modality (Figure 3B). Expression features were moderately more influential overall, with 77% of miRNAs showing higher weights for expression than for APA. This bias, however, was rather modest than extreme, and no outliers exhibited complete dependence on one feature type, consistent with previous observations.

From the gene perspective, we assessed whether target genes contributed systematically through expression or APA features. Among 8260 target genes across all HP-miRNAs, 70% exhibited a bias toward expression-derived contributions. Specific examples included CITED1, SLC52A2, and ACTG2, which were primarily expression-driven, whereas IFITM1 and PRDX6 were dominated by APA. Nonetheless, most genes featured in many miRNA models (>200) showed no strong preference, again highlighting the balanced contributions of both modalities (Figure 3C).

To further dissect modality-specific contributions, we stratified genes into APA- and expression-dominant groups based on the 10th and 90th percentile cutoffs of their dominance fraction. This classification yielded 891 APA-dominant and 768 expression-dominant genes across all HP-miRNAs. Notably, genes with high prevalence across multiple miRNAs typically exhibited only moderate dominance (Figure 3D).

When comparing variability across modalities, expression-dominant genes showed higher variance in both expression and APA relative to APA-dominant genes. For expression values, median coefficients of variation were 0.716 versus 0.650, and for APA, 0.225 versus 0.221. A Wilcoxon rank-sum test confirmed significantly greater variability in expression (p < 0.001) and APA (p < 0.01) for expression-dominant genes. Importantly, APA-dominant genes did not exhibit elevated APA variability across miRNAs, indicating that their predictive contribution reflects systematic APA regulation rather than noise (Figure 3E). Similarly, the top 100 recurrently used genes were rarely exclusive to APA or expression, but instead reflected mixed contributions (Figure 3F).

### Bimodal sign patterns reveal distinct Expression–APA relationships

Beyond their relative magnitudes, the coefficient signs for expression- and APA-derived features reveal how these two modalities tend to co-vary within our models. In general, positive coefficients for both expression and APA of target genes indicate that higher expression together with more distal 3′UTR usage is statistically associated with higher predicted miRNA levels, whereas negative coefficients for both modalities indicate the opposite—lower expression combined with more proximal site usage is statstically associated with lower predicted miRNA levels. These associations reflect predictive relationships and do not imply a specific direction of causality.

To assess whether individual miRNAs exhibit systematic patterns in how expression- and APA-derived contributions relate across their target genes, we summarized the distribution of coefficient sign combinations separately for each miRNA. miRNAs were then stratified according to whether concordant (++ and −−) or discordant (+− and −+) sign patterns predominated among their targets. This grouping was introduced to distinguish miRNAs for which expression and 3′UTR architecture tend to act in the same direction from those in which the two modalities contribute in opposing directions.

Applying this stratification revealed two dominant groups of miRNAs. About 423 HP-miRNAs (52 %) were dominated by the concordant sign patterns (++ and −−), in which expression and APA coefficients share the same sign. The remaining 390 HP-miRNAs (48 %) were dominated by the discordant sign patterns (+− and −+), in which coefficients have opposite signs (Figure 3G).

This bimodality highlights two prevalent modes by which expression and APA features relate to miRNA levels. Interestingly, these two groups also differed in predictive performance, with mean R^2^ values of 0.74 and 0.65, respectively (Wilcoxon rank-sum test, p < 0.01; Figure 3H). While the coefficients come from regularised models and cannot be interpreted as direct effect sizes, the systematic separation is consistent with opposing mechanisms such as compensatory biogenesis versus target-directed decay. However, because miRNA–target interactions are intrinsically bidirectional, increased target expression and 3′UTR lengthening may coincide with either higher or lower mature miRNA abundance, and the present framework cannot distinguish whether observed associations reflect dominant target-mediated sequestration, miRNA-driven repression, or a combination of both. Importantly, this does not diminish the biological relevance of the observed patterns, as the reproducible contribution of APA and expression features demonstrates that dynamic changes in target-site availability systematically shape steady-state miRNA levels.

### Global APA regulation as a determinant of miRNA logFC

To examine how APA modulates miRNA expression dynamics, we analyzed four perturbation experiments in which key APA-regulatory proteins were knocked down and compared with matched controls. These datasets included knockdowns of CFIm25 (two independent experiments), CFIm68, and CPSF6, factors that shape 3′UTR processing and thereby influence miRNA binding-site availability [24]. For each dataset we applied the MIRNAPEX pipeline to predict miRNA log-fold changes based solely on the observed gene-expression changes and APA shifts of their target genes.

Across all four perturbation experiments (CFIm25-KD-1, CFIm25-KD-2, CFIm68-KD, CPSF6-KD) we observed predicted miRNA logFC in both directions, with many exceeding an absolute value of 1 (Figure 4A). In CFIm25-KD-1 (4 miRNAs up-regulated and 15 down-regulated), CFIm25-KD-2 (6 up and 11 down), CFIm68-KD (20 up and 25 down) and CPSF6-KD (6 up and 14 down), the MIRNAPEX predictions indicated a range of miRNA logFC rather than a uniform shift. Notably, hsa-miR-182-5p showed consistent down-regulation in three of the four experiments.

**Figure 4.**
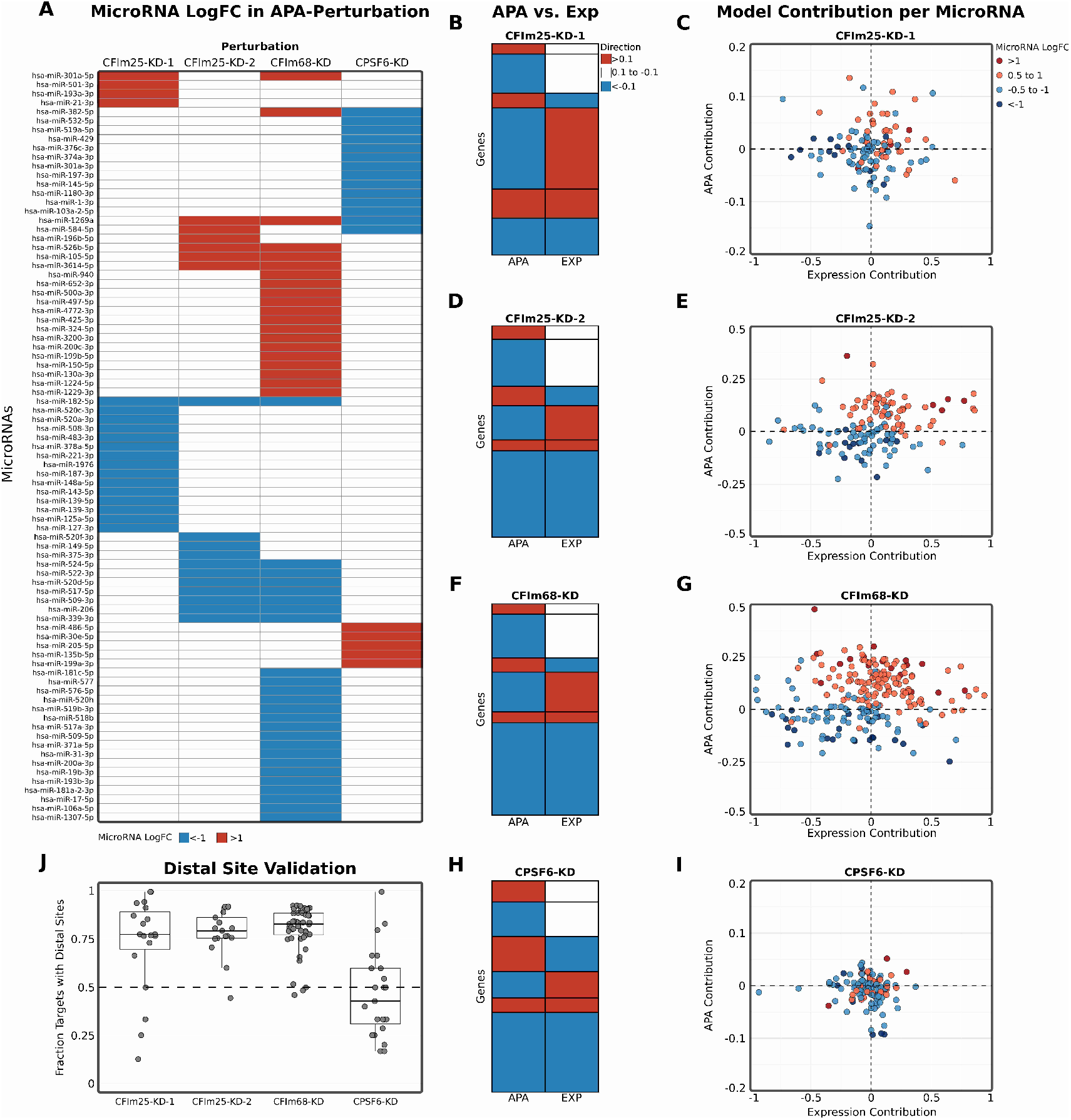
MiRNA behaviour in APA perturbation experiments. (**A**) Heatmap of MIRNAPEX-computed miRNA logFC across the four knockdown (KD) experiments of APA-regulatory factors (CFIm25-KD-1, CFIm25-KD-2, CFIm68-KD, CPSF6-KD). Only miRNAs with |logFC|>1 are displayed. (**B, D, F, H**) Heatmaps display, for each gene with an APA change of |ΔPDUI| > 0.1, the direction of 3′UTR change (APA column) together with the corresponding gene logFC (EXP column). Lengthening of 3′UTRs or positive gene logFC values are indicated in red, shortening or negative gene logFC values in blue, and no expression change in white. This representation highlights the directionality of APA changes relative to gene expression, showing whether 3′UTR lengthening or shortening coincides with increases or decreases in gene logFC across APA perturbation experiments. (**C, E, G, I**) Scatter plots show, for each miRNA across the perturbation experiments, the combined (unscaled) contribution of gene expression changes versus APA changes to the predicted miRNA logFC as computed by MIRNAPEX. Points are shaded according to the predicted miRNA logFC values across the defined thresholds. (**J**) Boxplots showing the proportion of APA-changed target genes that harbour at least one distal binding site for the same miRNA (|logFC|>1) in each perturbation experiment.

Since the direction of individual miRNA changes correlated with the expression and APA shifts of their target genes, we investigated how many genes with APA changes also display corresponding differences in gene expression, and how the direction of 3′UTR change differences relates to the gene logFC to reveal the directionality of these effects.

Across the four perturbation experiments, we observed widespread changes in 3′UTR usage, with a clear predominance of shortening events and varying degrees of buffering, expression changes in the opposite direction to the APA effect, likely reflecting compensatory mechanisms such as altered miRNA activity as consequence of binding site modulation.

In CFIm25-KD-1, 6721 genes displayed altered 3′UTR usage (defined as ΔPDUIs ≥ 0.05 for genes with |logFC| ≥ 0.1), with 5325 (79 %) showing shortening; about 1698 (25 %) of these APA-changed genes exhibited opposite (buffering) expression shifts (Figure 4B). In CFIm25-KD-2, 385 genes showed altered 3′UTR usage, with 286 (74 %) showing shortening and 175 (46 %) displaying opposite expression changes (Figure 4D). In CFIm68-KD, 6902 genes had altered 3′UTR usage, with 5742 (83 %) showing shortening and roughly 1767 (26 %) exhibiting opposite expression changes (Figure 4F). In CPSF6-KD, 361 genes displayed altered 3′UTR usage, with 240 (67 %) showing shortening and about 105 (29 %) showing opposite expression changes (Figure 4H).

We tested whether APA remodeling translates into predictable shifts in miRNA levels by splitting each miRNA’s predicted change into two additive components: one reflecting expression of target genes and the other reflecting APA patterns, keeping unscaled values to highlight the direction and relative magnitude.

We found that miRNAs with larger predicted changes cluster in quadrants where the APA component change and the observed miRNA change point in the same direction. In CFIm25-KD-1, CFIm25-KD-2, and CFIm68-KD this concordance is significant (one-sided Fisher’s exact test, p < 0.01 for miRNAs with |logFC| > 0.5), indicating that a stronger APA contribution is associated with larger miRNA shifts and vice versa (Figure 4C;E;G). In CPSF6-KD no enrichment is observed (p = 0.53), consistent with this perturbation showing the lowest fraction of 3′UTR-shortened genes among the datasets considered Figure 4I). This pattern further emphasizes that extensive gene shortening in APA perturbations coincides with the largest shifts in miRNA levels and that, where shortening is limited, the APA component contributes less strongly to miRNA log fold-changes.

For each miRNA with a predicted change, we assessed whether the 3′UTR shortening or lengthening of its target genes in the same experiment is associated with altered availability of binding sites for that specific miRNA. Using the microT predictions, APA-changed genes were screened for the presence of at least one binding site for the same miRNA in the region between the proximal and distal polyadenylation sites. This analysis showed that in CFIm25-KD-1, CFIm25-KD-2 and CFIm68-KD a substantial fraction of shortened targets indeed contained a distal binding site for the same miRNA, with median values of 77.8%, 79.7% and 83.1% of APA-changed genes, respectively. In CPSF6-KD the median proportion was much lower (42.9%), consistent with the weaker shortening seen in this dataset (Figure 4J). These results validate that the predicted miRNA shifts reflect real changes in binding-site dosage caused by APA remodeling. To further validate PDUI as a proxy for miRNA binding-site availability, we extended this analysis to all expressed miRNAs in each perturbation dataset, independent of their predicted logFC. Across datasets, APA-regulated target genes frequently harbored distal binding sites for the corresponding miRNA, with median fractions of 72.7% in CFIm25 KD—HCT116, 70.7% in CFIm25 KD—HEK293, 70.7% in CFIm68 KD—HEK293, and 56.5% in CPSF6 KD—HEP3B. These genome-wide results support PDUI as a meaningful measure of miRNA binding-site dosage in the present analyses.

In summary, our findings show a strong statistical concordance between APA-driven target shortening and miRNA logFC. This suggests that at least part of the apparent miRNA DE we observe may reflect changes in binding-site availability rather than direct changes in miRNA transcription.

### APA-driven loss of TDMD trigger sites coincides with miRNA abundance shifts

As APA-perturbation experiments globally shorten 3′UTRs, we asked whether this remodeling also affects established TDMD interactions. We therefore investigated eight curated trigger–miRNA pairs, as well as one negative control, in which the trigger transcript harbours highly complementary sites known to direct miRNA decay and for which we observed APA changes of the trigger gene. These included CYRANO with hsa-miR-7-5p, SDC2 with hsa-miR-15a-5p, SERTAD3 with hsa-miR-92a-3p, SSR1 with hsa-miR-218-5p, TRIM9 with hsa-miR-218-5p, TDP1 with hsa-miR-320a-3p, NREP with hsa-miR-29b-3p, and BCL2L11 with hsa-miR-221-3p, alongside BCL2L11 with hsa-miR-221-5p as a negative control [14,46,47].

For each APA-perturbation dataset, we extracted the trigger genes, quantified their ΔPDUI values to assess 3′UTR remodeling, and compared these changes with MIRNAPEX-predicted log fold-changes of the corresponding mature miRNAs (Figure 5A–I). This analysis directly tests whether loss of distal 3′UTR regions harbouring TDMD trigger sites under APA perturbation is associated with increased abundance of the targeted miRNAs.

**Figure 5.**
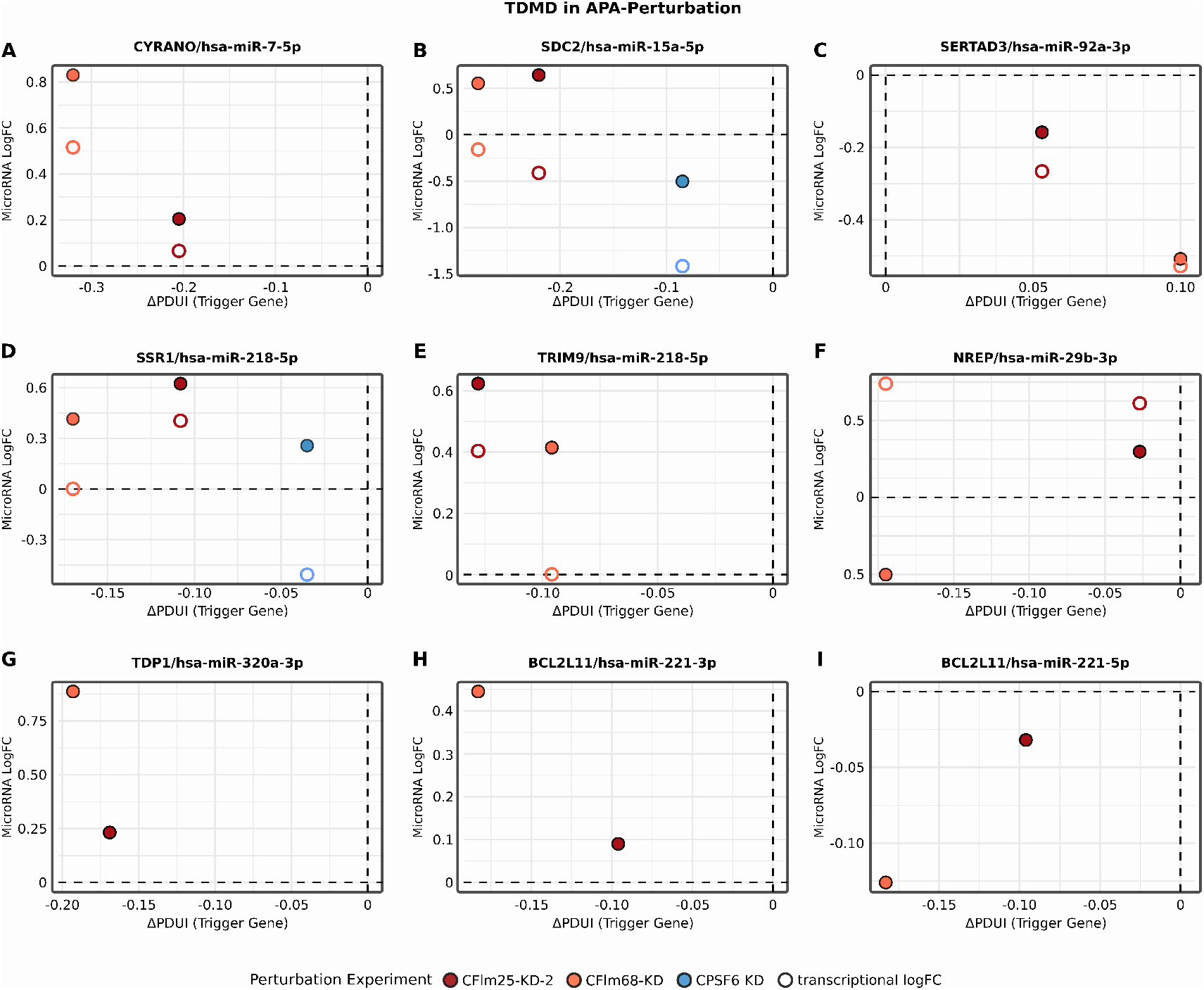
APA-driven 3′UTR shortening of TDMD trigger genes and miRNA abundance changes. Scatter plots relating APA changes of TDMD trigger genes to the log fold-change of their paired mature miRNAs across APA-perturbation datasets. Each panel corresponds to one trigger–miRNA pair: **(A)** OIP5-AS1 with hsa-miR-7-5p, **(B)** SDC2 with hsa-miR-15a-5p, **(C)** SERTAD3 with hsa-miR-92a-3p, **(D)** SSR1 with hsa-miR-218-5p, **(E)** TRIM9 with hsa-miR-218-5p, **(F)** NREP with hsa-miR-29b-3p, **(G)** TDP1 with hsa-miR-320a-3p, **(H)** BCL2L11 with hsa-miR-221-3p, and **(I)** BCL2L11 with hsa-miR-221-5p (negative control). Filled symbols indicate mature miRNA logFC values predicted by MIRNAPEX, while open symbols represent transcriptional proxy logFC estimates. Points are coloured by perturbation dataset. The x-axis shows ΔPDUI values (KD − control), with negative values indicating 3′UTR shortening, and the y-axis shows log fold-changes.

Across the majority of TDMD pairs, negative ΔPDUI values of the trigger gene, indicative of 3′UTR shortening, coincided with increased mature miRNA abundance. This trend was particularly evident for pairs involving CYRANO with hsa-miR-7-5p (Figure 5A), SDC2 with hsa-miR-15a-5p (Figure 5B), SSR1 with hsa-miR-218-5p (Figure 5D), TRIM9 with hsa-miR-218-5p (Figure 5E), TDP1 with hsa-miR-320a-3p (Figure 5G), and BCL2L11 with hsa-miR-221-3p (Figure 5H). In these cases, the strongest miRNA up-regulation was observed in datasets exhibiting pronounced trigger 3′UTR shortening, consistent with reduced TDMD-mediated degradation following loss of distal trigger regions **[16]**. By contrast, SERTAD3 with hsa-miR-92a-3p (Figure 5C) did not exhibit consistent 3′UTR shortening across perturbations and accordingly showed decreased miRNA abundance in conditions associated with 3′UTR lengthening, supporting the directional relationship between trigger 3′UTR architecture and miRNA stability. NREP with hsa-miR-29b-3p (Figure 5F) deviated from the general trend, displaying divergent miRNA responses across perturbations despite trigger shortening, suggesting that additional regulatory inputs or context-dependent effects may modulate TDMD efficiency for this pair.

Importantly, the negative control pair BCL2L11 with hsa-miR-221-5p (Figure 5I) did not show systematic miRNA up-regulation despite APA changes of the trigger gene, indicating that TDMD sensitivity is arm-specific and reinforcing that the observed effects are not a general consequence of trigger gene expression changes.

To distinguish post-transcriptional effects from altered miRNA production, we additionally compared mature miRNA logFC values with transcriptional proxy measurements derived from host-gene or pri-miRNA-associated expression estimates (hollow circles in Figure 5A–F). For all TDMD pairs showing increased mature miRNA abundance under trigger shortening, the transcriptional proxy logFC was lower or unchanged, indicating that the observed miRNA up-regulation cannot be explained by increased transcription. In contrast, NREP with hsa-miR-29b-3p showed no consistent separation between transcriptional proxy and mature miRNA changes, in line with its context-dependent behaviour.

Taken together, these results indicate that MIRNAPEX captures TDMD-linked miRNA behaviour in the majority of curated cases, and that APA-driven loss of distal trigger regions is frequently associated with increased mature miRNA abundance independent of transcriptional changes. This supports the view that a substantial fraction of miRNA expression changes observed under APA perturbation reflects altered TDMD site availability rather than solely changes in miRNA transcription.

## Discussion

Steady-state miRNA levels arise from a dynamic balance of biogenesis, target engagement and decay, including TDMD, yet DE of miRNAs is often interpreted as evidence of altered post-transcriptional regulation [11,12,16]. Within this balance, APA reshapes 3′UTR isoform usage and therefore affects effective dosage of canonical binding sites and highly complementary decay triggers [24,48]. To test whether such transcriptome remodeling predicts miRNA logFCs between conditions, we developed MIRNAPEX, which reads out target-centric features derived jointly from gene expression changes and APA.

Across miRNAs, features derived solely from APA were predictive on their own and, when combined with mRNA expression, consistently improved performance, showing that 3′UTR remodeling contributes information absent from total transcript levels. Mechanistically this is consistent with the idea that site dosage and binding strength together determine AGO occupancy and repression efficacy [49–51].

From a gene-centric perspective, many targets contributed to prediction mainly through expression features, reflecting changes in total abundance and baseline miRNA binding site load, while others were dominated by APA, consistent with isoform switches that add or remove distal sites or decay triggers without large changes in total transcript levels [24].

Both target-gene expression and APA influence predicted miRNA logFC: expression dominates overall, but APA leads for many targets. Interestingly, miRNAs fall into two behavior classes: in one, expression and APA effects align, more transcript and longer 3′UTRs go with higher miRNA levels, and in the other, they oppose, so increased site availability is associated with lower miRNA. This bimodal pattern suggests distinct regulatory modes for different miRNAs, not just variation in target abundance; such modes are consistent with competition and sequestration effects and observed differences in miRNA-mRNA network behavior in recent studies [52].

Applying MIRNAPEX to experimental data where core APA regulators were knocked down revealed broad 3′UTR shortening, consistent with the known global impact of APA perturbations, with variable buffering at the expression level and predicted miRNA changes in both directions [28,53,54]. Decomposing predictions showed that the largest miRNA shifts clustered where the APA contribution and miRNA direction agreed, especially in perturbations that induce extensive shortening, whereas when shortening was limited, the contribution was weaker. Screening shortened targets confirmed that a substantial majority harbored distal sites for the same miRNA in these datasets, validating that MIRNAPEX effects stemming from site dosage rather than spurious correlations.

For TDMD trigger–miRNA pairs such as BCL2L11–miR-221-3p, NREP–miR-29b-3p, SSR1–miR-15a-5p, TDP1–miR-320a-3p and TRIM9–miR-218-5p, 3’UTR shortening generally coincided with higher predicted miRNA abundance, consistent with relief from decay [14,16]. TDMD triggers located in the 3′UTR have been shown to degrade miRNAs more effectively than identical triggers placed in coding sequences [46], which supports our focus on 3′UTR site loss in interpreting miRNA changes. A recent study found an endogenous TDMD trigger with minimal non-canonical 3′-end base-pairing that is nevertheless sufficient to induce degradation of the miR-279 family [55]. This suggests that many APA-associated miRNA expression changes could reflect widespread but previously uncharacterized TDMD triggers captured by MIRNAPEX. These findings further indicate that MIRNAPEX is sensitive to mechanistically defined decay events embedded within broader APA remodeling.

MIRNAPEX has some important limitations. It relies on bulk RNA-seq–derived APA metrics and selection of features based on predicted target sets, which are imperfect proxies for the true binding landscape. Bulk data also mix cell types and states, so shifts in composition could confound expression and 3′UTR usage. Furthermore, coefficients from regularized models help interpretation but don’t explain causal effect sizes and cannot fully separate biogenesis from decay. Finally, curated TDMD interactions are incomplete and context-dependent, limiting ground-truth validation.

Integrating direct AGO-binding data with isoform-resolved APA profiles and more detailed predictors of site efficacy will sharpen our understanding of how target landscapes influence miRNA levels. Applying such analyses at single-cell or time-course resolution could help separate cell-state effects from true regulatory changes, while controlled perturbations with matched mRNA, APA and small-RNA measurements would provide stricter benchmarks for testing mechanistic hypotheses.

Our findings suggest that a more comprehensive view of miRNA regulation can be obtained when dynamic changes in target-site availability and decay processes are explicitly taken into account. Incorporating these dimensions has the potential to improve the quality of commonly applied analysis workflows, strengthen functional interpretations and increase the reliability of biomarker discovery.

## Supplementary Data

**Supplementary Figure 1:**
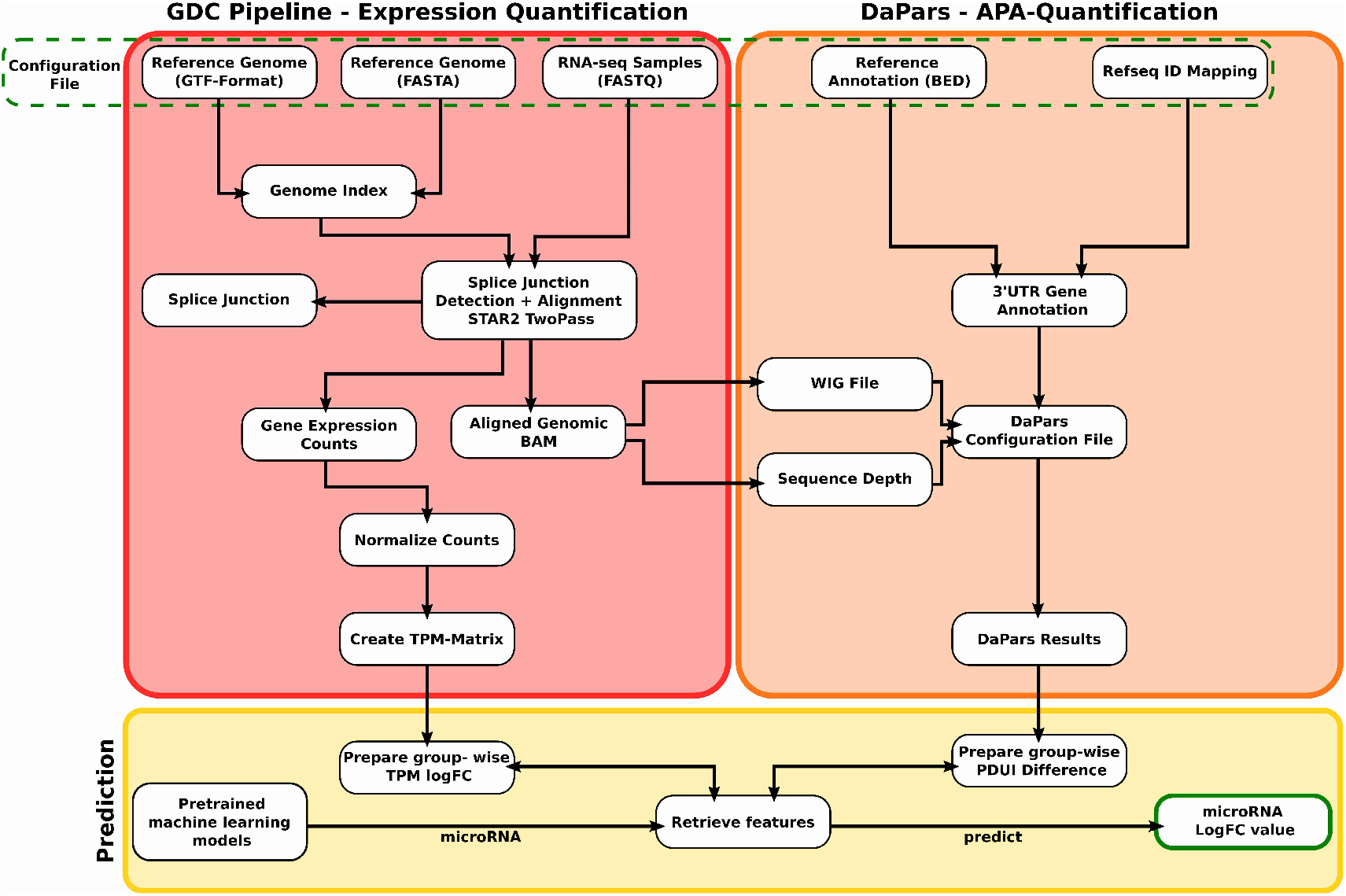
Illustration of the MIRNAPEX workflow.

Supplementary Table 1: Benchmarking metrics for the pretrained miRNA models using a 1000-feature set with EN.

## Acknowledgements

Parts of this research were conducted using the supercomputer MOGON 2 and/or advisory services offered by Johannes Gutenberg University Mainz (hpc.uni-mainz.de), which is a member of the AHRP (Alliance for High Performance Computing in Rhineland Palatinate, www.ahrp.info) and the Gauss Alliance e.V..

## Author contributions

Conceptualization, M.A.A.-N and M.C.; Data curation, M.C.; Formal analysis, M.C.; Investigation, M.C.; Methodology, M.A.A.-N, M.C., P.M., M.S. and F.M.; Supervision, M.A.A.-N; Visualization, M.C.; Writing – original draft, M.C.; All authors reviewed and edited the manuscript. All authors have read and agreed to the published version of the manuscript.

## Data and code availability

The MIRNAPEX workflow is openly available at https://github.com/mcihan0bioinf/MIRNAPEX and archived on Zenodo (https://doi.org/10.5281/zenodo.17474139). It is provided as a fully defined computational pipeline within a conda environment. Model coefficients and training codes are accessible through this repository.

## Statements & Declarations

### Funding

This work was supported by the Deutsche Forschungsgemeinschaft (DFG, German Research Foundation) Project number 318346496 - SFB1292/2 TP19N (to FM). No additional funding was received for the preparation of this manuscript.

### Competing Interests

The authors have no relevant financial or non-financial interests to disclose.

### Ethics approval

This study did not involve any new experiments on human participants or animals. All data were obtained from public databases and therefore no ethics approval was necessary.

## Notes

https://github.com/mcihan0bioinf/MIRNAPEX

### Competing Interest Statement

The authors have declared no competing interest.

### Summary of Updates

Figure 2 revised; Figure 5 revised; expanded TDMD validation with additional trigger-miRNA pairs and a negative control; extended analysis of transcriptional expression proxies to distinguish transcriptional and post-transcriptional effects; cross-validation of results using MirGeneDB-annotated microRNAs.

