## Supplementary Figure 1 for "Target-site Dynamics Explain a Large Share of Apparent MicroRNA Differential Expression"

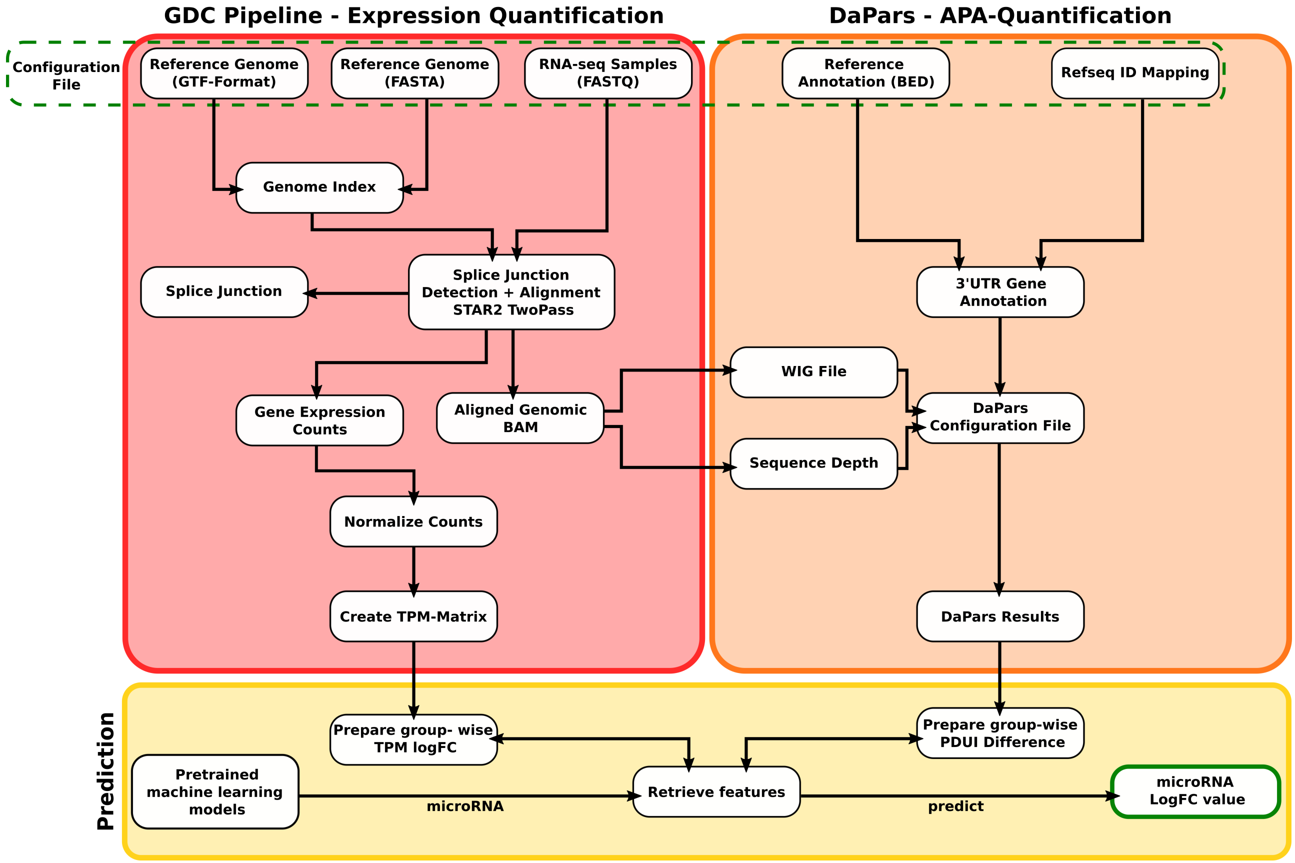


**Supplementary Figure 1. MIRNAPEX workflow.**
Schematic of the MIRNAPEX pipeline, showing how RNA-seq reads are aligned and quantified, 3′UTR usage is estimated with DaPars, and expression and APA features are combined with reference annotations to feed the pre-trained models that predict miRNA log-fold changes.
